# 1014F-*kdr* and G119S *ace-1* mutations conferring insecticide resistance detected in the *Culex pipiens* complex in Morocco

**DOI:** 10.1101/2023.08.14.553296

**Authors:** Souhail Aboulfadl, Chafika Faraj, Karim Rabeh, Btissam Ameur, Bouchra Belkadi, Fouad Mellouki

## Abstract

*Culex pipiens* mosquitoes are competent vectors for several arboviruses worldwide. In Morocco, they specially transmit West Nile virus (VWN) and Rift Valley fever virus (VFVR) (Bkhache et al., 2018). The repeated treatment with chemical insecticides in areas highly exposed to the mosquitoes’ nuisance leads to the resistance development. The lack of data on resistance mechanisms in Morocco allow us to evaluate both of the levels of DDT resistance accompanied by the frequency of the mutated gene associated L1014F *kdr*, and the levels of bendiocarb resistance accompanied by the frequency of the mutated gene associated G119S *ace-1*^*R*^ respectively in both regions Benslimane and Mohammedia.

**Methods:** Mosquito larvae were collected from two different breeding sites: El Jazeera from Benslimane region and Ouled Hamimoun from Mohammedia region. Adults were reared from collected immature stages in the laboratory at 28 ± 1 °C with 80% relative humidity. Standard WHO insecticide susceptibility tests were conducted on adult females emerged from collected larvae. Specimens were identified as *Culex pipiens* following morphological criteria under microscope using Culicidae of African Mediterranean software (Giner et al., 1999). Identified mosquitoes were then tested for the presence of the L1014F *kdr* and G119S *ace-1*^*R*^ mutations using PCR assay.

**Results:** Our results showed that all the tested populations have a resistance to DDT 4% with a mortality rate ranging from 2% to 74%, and to bendiocarb 0.1% with a mortality rate ranging from 32% to 74%. In plus, 30 individuals from each region were screened for both mutations L1014F *kdr* and G119S *ace-1R*. The molecular identification of survivor genes shows the presence of L1014F *kdr* mutation in both region with different frequencies. The allelic frequency was low with 0.15% in Mohammedia region and complete absence of resistant homozygotes alleles 1014F/1014F in the same region, but average with 0.65% in Benslimane region. However, the G119S *ace-1*^*R*^ mutation was absent in both regions. These results are consistent with those published by Aboulfadl et al. (2020) on the same populations and who found that the resistance of Mohammedia populations is completely metabolic and based only on cytochrome P450 enzymes while that of Benslimane populations is partially metabolic.

**Conclusion:** The study highlights the implication of L1014F *kdr* mutation in DDT resistance and G119S *ace-1*^*R*^ mutation in bendiocarb resistance in *Cx. pipiens* populations from two regions in Morocco.

## Introduction

Vector born diseases transmitted by mosquitoes are in a major concern worldwide. Mosquitoes remain important vectors by inducing 1M death and over than 700M infections worldwide every year. *Culex pipiens*, in particular, is a vector of lymphatic filariasis that threaten 1.1 billion of people worldwide (WHO, 2016). In Morocco, *Cx. pipiens* is a competent vector of several arboviruses such as West Nil virus and Rift Valley Fever virus (Amraoui et al. 2014). It has been suspected as vector of West Nil epizootics in 1996 which caused 94 equine cases including 42 dead horses and one human case in Benslimane, in 2003 in Kenitra and in 2010 in Mohammedia (Schuffenecker et al. 2005).

In the absence of effective vaccines, the mosquito control remains the unique way to limit pathogen transmission. Thus, the use of neurotoxic insecticides plays a major role in the prevention and control of vector-borne diseases. However, the frequent use of insecticides (mainly pyrethroids and organophosphates) has contributed to select several resistance mechanisms in targeted mosquito populations. In Morocco, most of studies showed significant resistance of *Cx. pipiens* to all chemical insecticides: organophosphates in *Culex. pipiens* larvae from Rabat, Sale, Skhirate, Mohammedia (Faraj et al. 2002), Smir (El Joubari et al. 2015), Fes (El Ouali Lalami et al. 2014), and Mohammedia, Benslimane, Skhirate and sale (Aboulfadl et al. 2020); organochlorines in *Cx. pipiens* larvae from Khemisset (Larhbali et al. 2010); pyrethroids in *Cx. pipiens* larvae from Casablanca, Tanger, and Marrakech (Bkhache et al. 2016). The resistance occurred in almost all regions in central Morocco needs to be well studied for a better understanding to the mechanisms implicated by the mosquito. In plus, the operational impact of a given resistance might not be achieved yet what makes the investigation of the resistance intensity primordial.

The objective of our study was to investigate the resistance mechanisms related to target site modifications, especially G119S *ace-1*^*R*^ (Gly119 replaced by Ser) in the acetylcholinesterase enzyme coded by *ace-1* gene target site of carbamates and organophosphates; and leucine to phenylalanine L1014F-*kdr* in the voltage gated sodium channel (vgsc) coded by *kdr* gene target site of organochlorines and pyrethroids. Our results give a better understanding to molecular mechanisms implicated by mosquitoes against insecticides, and thus can be used as a guide to the choice of mosquito control and resistance management strategy.

## Materials and methods

### Ethics statement

No specific authorization is required for activities in the field that do not concern endangered or protected species. This work was collaborated with the National Institute of Hygiene in Morocco which is a public health and scientific research institution under the supervision of the Ministry of Health. In this context, it participates in vector control activities that allow it to operate without specific authorization to access breeding sites and mosquito collections.

### Mosquito collection and rearing

Larvae of *Cx. pipiens* were collected from March to Mai 2020 in two urban districts: Ouled Hamimoun (33.673748, –7.445547) from Mohammedia, and El-Jazeera (33.780420, – 7.235220) from Benslimane. Larvae and pupae were then brought for adult emergence to National Institute of Hygiene Laboratory of Rabat. Adults were maintained in cages at 27 ± 2°C, 80% of humidity, 12h:12h photoperiod and supplied with a 10% of sugar solution. The F1 mosquitoes were used for all insecticide susceptibility tests and molecular biology investigations.

### Adult bioassays

Adult bioassays were conducted following adapted WHO recommendations (WHO, 2016). We tested adult susceptibility using WHO discriminating dosages (DD) of two insecticides belonging to two chemical classes: organochlorine (DDT 4%) and carbamate (bendiocarb 0.1%). Four batches of 25 *Cx. pipiens* unfed females, aged 2-5 days, were exposed for 4h to DDT insecticide-treated papers, 2h to bendiocarb. The number of mosquitoes knocked down while were exposed to DDT was recorded at intervals of 30 min. After the exposure period, all mosquitoes were transferred to observation tubes with 10% sugar solution supplement. Batches exposed to oil-treated papers were run in parallel and used as control. Knock-down time (KdT50 and KdT90 indicating 50% and 90% of knocked-down tested populations respectively) were calculated using WinDL32 software (Giner et al., 1999). Mortality rate was recorded after 24h. Dead and alive mosquitoes were kept in tubes with silica gel. All preserved mosquitoes were stored in a freezer at -20 °C for resistant gene screening.

### Genotyping of target-site mutations

Genomic DNA was extracted from 30 *Cx. pipiens* adult females from each of the two populations studied of Mohammedia and Benslimane using TNES technique. The extracted DNA was used to genotype L1014F *kdr* mutation of the VGSC and G119S *ace-1* mutation. The L1014F was genotyped using the double PCR-based assay described by Martinez-Torrez et al. (1999) with slight modification following PCR conditions: 1 min at 95ºC followed by 35 cycles of 94ºC for 30 sec, 50ºC for 30 sec in 1^st^ PCR (59°C for 30 sec in 2^nd^ PCR) and 72ºC for 30 sec, with a final extension of 5min at 72ºC. The DNA fragments were visualized by electrophoresis on 1.5% agarose gel with ethidium bromide and observed under ultraviolet light. The G119S *ace-1* mutation was investigated using the PCR-RFLP (PCR Restriction Fragment Length Polymorphism) test developed by Weill et al. (2004). The PCR conditions used were as follows: 5 min at 94ºC, 30 cycles (94ºC for 30 sec, 47ºC for 30 sec and 72ºC for 1 min) and a final extension of 5 min at 72ºC. PCR products were digested with the restriction enzyme AluI for the ace-1 gene, at 37°C for 3h according to the manufacturer’s instructions. The digestion products were subjected to electrophoresis on 2% agarose gel colored with ethidium bromide and observed under UV light. Primers used are shown in Table 1.

**Table1:**
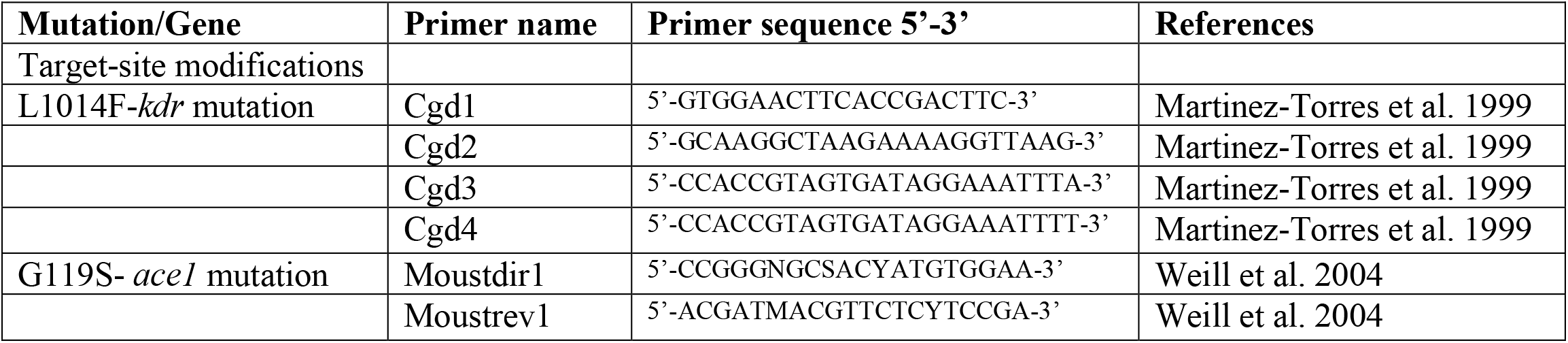
Primers used for the screening of target-site modifications.

### *DNA* diagnostic test for *kdr* alleles in single mosquitoes

150ng of genomic DNA prepared as above were combined in a 25 ml total volume with the four primers Cgd1, Cgd2, Cgd3 and Cgd4. 1^st^ PCR was to detect the substitution of leucine to phenylalanine: 1.25 μl [0.5μM] Cgd1 + 2.5 μl [1μM] Cgd2 + 1.25 μl [0.5μM] Cgd3 + 5 μl buffer reaction + 0.5 Taq DNA. The 2^nd^ PCR was to detect the wild allele: 1.25 μl [0.5μM] Cgd1 + 2.5 μl [1μM] Cgd2 + 1.25 μl [0.5μM] Cgd4 + 5 μl buffer reaction + 0.5 Taq DNA. The 1^st^ PCR reaction conditions were 1 min at 95°C, 30s at 94°C, 30s at 50°C and 30s at 72°C for 35 cycles with external extension step at 72°C for 5 min. The 2^nd^ PCR reaction conditions were 1 min at 95°C, 30s at 94°C, 30s at 59°C and 30s at 72°C for 35 cycles with external extension step at 72°C for 5 min. Amplified fragments were analyzed by electrophoresis on a 1.5% agarose gel and were visualized by ethidium bromide staining under UV light.

### Statistical analysis

Following exposure of *Cx. pipiens* populations to insecticides, those with recorded mortalities between 98–100% were rated as susceptible. Populations showing mortality below 98% were rated as suspicious of resistance and populations showing mortality below 90% were rated as resistant (WHO, 2004). Genotype Allelic frequencies were calculated using the following formula f(R)= (p + q)2 = p2 + 2pq + q2 = 1, where p2 is the genotype frequency of the homozygote susceptible allele AA, 2pq is the genotype frequency of the heterozygote resistance allele AB and q2 is the genotype frequency of the homozygote resistance allele BB.

## Results

### Insecticide resistance profile of *Cx. pipiens* for DDT and bendiocarb

The complex of *Culex pipiens* mosquitoes collected from the study area showed resistance to all tested insecticides (Table 1). For organochlorates (DDT) and carbamates (bendiocarb), resistance was observed in mosquitoes with mortality rates of 2% and 32%, respectively in Benslimane, and with mortality rates of 74% both in Mohammedia. In Benslimane, DDT caused only 2% of mortality, which revealed a low susceptibility, while in Mohammedia it caused only 74% of mortality. KDT50 and KDT90 for DDT were 166 min and 816 min respectively, in Mohammedia and 1771 min and 6978 min respectively, in Benslimane (Table 1).

**Table 1:** Susceptibility status of *Culex pipiens* from Mohammedia and Benslimane against DDT (OC) and bendiocarb (CX).

|  | Mohammedia |  |  |  |  | Benslimane |  |  |  |  |
| --- | --- | --- | --- | --- | --- | --- | --- | --- | --- | --- |
| Insecticide | N | KdT50 (min) | KdT90 (min) | %Mr | S | N | KdT50 (min) | KdT90 (min) | %Mr | S |
| **DDT 4%** | 91 | 166 | 816 | 74 | R | 100 | 1771 | 6978 | 2 | R |
| **Bendiocarb 0.1%** | 100 | - | - | 74 | R | 94 | - | - | 32 | R |

N: Number of mosquitoes; KdT50: Time (min) during which 50% of the tested population undergoes a Knock down; KdT90: Time during which 90% of the tested population undergoes a Knock down; %Mr: Percentage of mortality; R: Resistant (% Mr < 90)

### Genotyping of L1014F *kdr* and G119S *ace-1* mutations

A high frequency of the L1014F *kdr* mutation was detected in *Cx. pipiens* populations from Benslimane, while in Mohammedia, the frequency of L1014F *kdr* mutation was very low. The frequency of genotypes was represented in Table 2, 3. The frequency of the mutant and resistant allele (F) was higher than the wild susceptible allele (L): 0.65 for Benslimane, and lower than the wild susceptible allele (L) :0.15 for Mohammedia. All specimens of susceptible mosquitoes had 1014 L/1014 L genotype in both regions. Among the 15 resistant mosquitoes from Benslimane, 4 had 1014 F/1014 F genotype, 5 had 1014 F/1014 L genotype and 1 had 1014L/ 1014 L genotype (Table 2). In Mohammedia, from all resistant mosquitoes, no 1014 F/1014 F genotype was found, 3 had 1014 F/1014 L genotype and 7 had 1014 L/1014 L genotype (Table 3). The resistant homozygote genotype (F/F) was found only in the populations of Benslimane (40% of individuals) and completely absent in populations of Mohammedia (0%). The proportion of heterozygote genotype (L/F) found in Benslimane (50%) was higher than the proportion observed in mosquitoes from Mohammedia (30%). The wild homozygote susceptible genotype (L/L) was very high in the *Cx. pipiens* from Mohammedia (70% of individuals), and very low in populations from Benslimane (10% of individuals).

**Table 2:** Frequencies of *kdr* mutation according to the phenotypic status (resistant/susceptible) of *Cx. pipiens* in Mohammedia.

| Genotype (%) |  |  |  |  |  |  |  |  |
| --- | --- | --- | --- | --- | --- | --- | --- | --- |
| Region | Control |  | Phenotype | N | 1014L/1014L<br>*n* (%) | 1014L/1014F<br>*n* (%) | 1014F/1014F<br>*n* (%) | Frequency of<br>allele 1014F (%) |
|  | N | Dead |  |  |  |  |  |  |
| Mohammedia | 40 | 0 | Resistant | 10 | 7 (70) | 3 (30) | 0 | 0,15 |
|  |  |  | Susceptible | 5 | 5 (100) |  |  |  |

Abbreviation: N, number of individuals tested

**Table 3:**
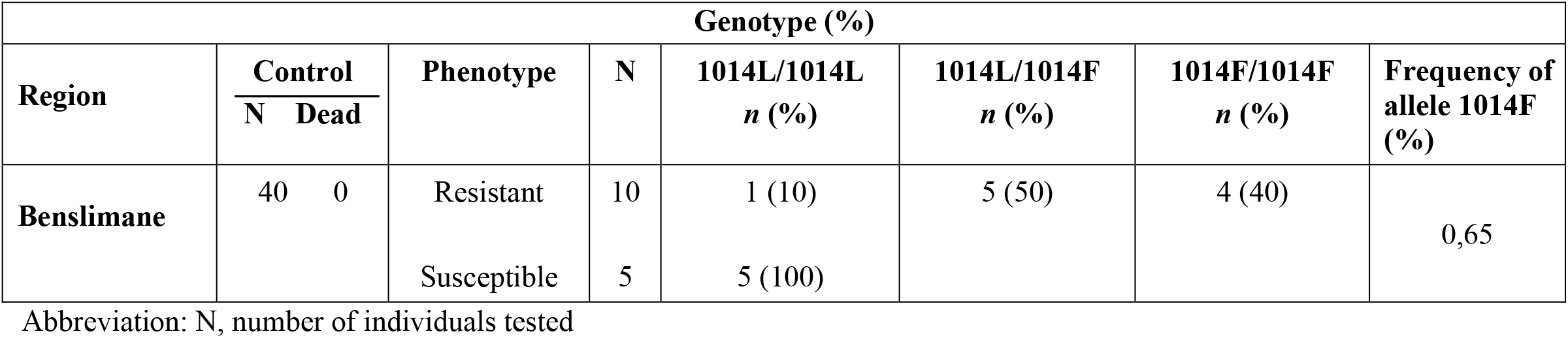
Frequencies of *kdr* mutation according to the phenotypic status (resistant/susceptible) of *Cx. pipiens* in Benslimane.

| Genotype (%) |  |  |  |  |  |  |  |
| --- | --- | --- | --- | --- | --- | --- | --- |
| Region | Control<br>N Dead | Phenotype | N | 1014L/1014L<br>*n* (%) | 1014L/1014F<br>*n* (%) | 1014F/1014F<br>*n* (%) | Frequency of<br>allele 1014F<br>(%) |
| Benslimane | 40 0 | Resistant | 10 | 1 (10) | 5 (50) | 4 (40) | 0,65 |
|  |  | Susceptible | 5 | 5 (100) |  |  |  |

Abbreviation: N, number of individuals tested

Regarding the G119S *ace-1* mutation, no *ace-1* mutation was found in any of the populations (Figure 2).

**Figure 1:**
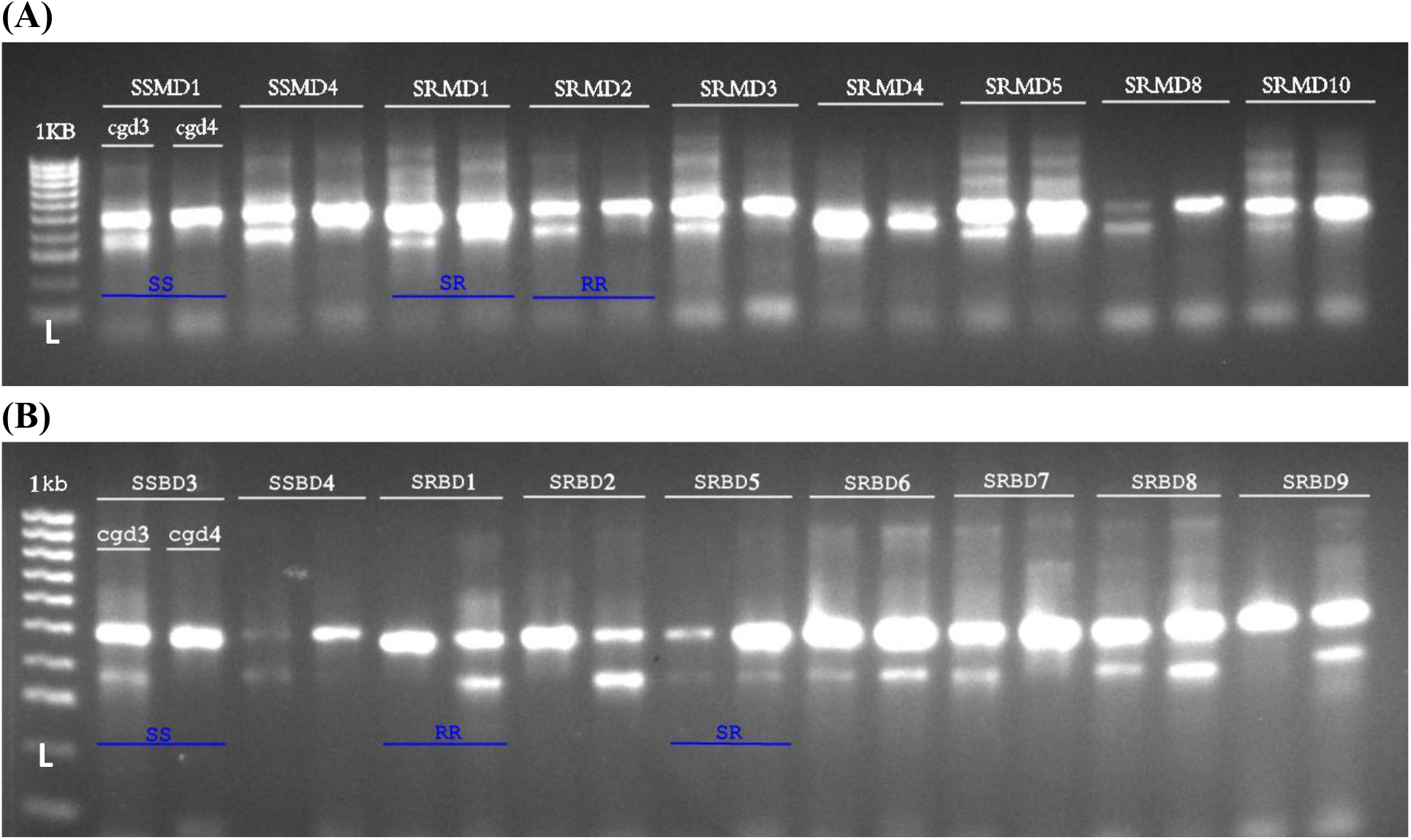
Detecting L1014F in single individuals. (A) PCR products obtained using double PCR-based assay on nine *Culex pipiens* strains from Mohammedia after separation on a 1.5% agarose gel. Lane L: 1 kb ladder; lanes 2-10: products obtained using genomic DNA of single mosquitoes as templates; lanes SSMD1 and SSMD4: susceptible insects; lanes SRMD(1,2,3,4,5,8,10): kdr insects. (B) PCR products obtained using double PCR-based assay on nine *Cx. pipiens* strains from Benslimane after separation on a 1.5% agarose gel. lanes SSBD3 and SSBD4: susceptible insects; SRBD(1,2,5,6,7,8,9): kdr insects. ss: susceptible homozygous mosquitoes; sr: resistant heterozygous mosquitoes; rr: homozygous resistant mosquitoes.

**Figure 2:**
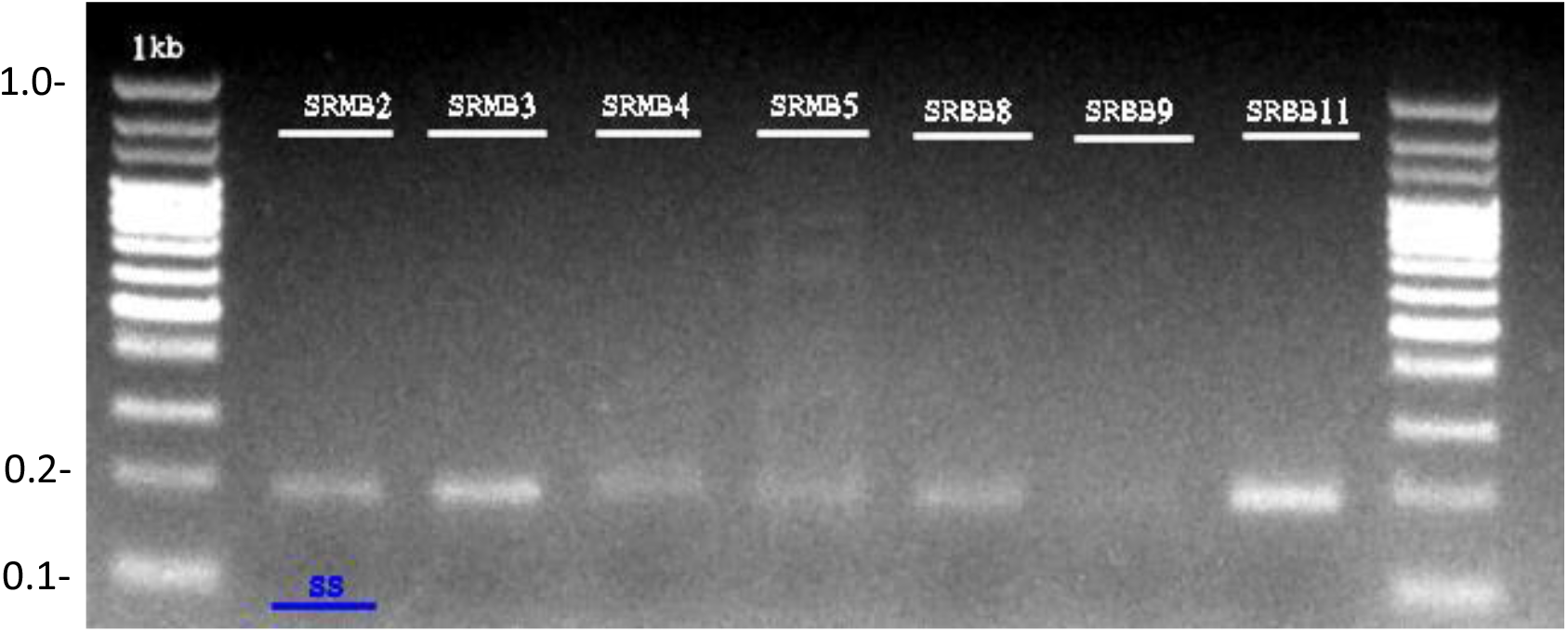
Detecting G119S in single individuals. Diagnostic PCR-RFLP test to detect the G119S mutation in *Cx. pipiens* individual mosquitoes. Genomic DNA amplification with the degenerated primers Moustdir1 and Moustrev1 produced a 194 bp fragment, which is undigested by AluI for susceptible homozygous mosquitoes (SS).

## Discussion

This paper is the continuous study published by Aboulfadl et al. (2022) in evaluating the resistance intensity and the metabolic mechanisms involved in the disease vector *Culex pipiens* from Mohammedia, Benslimane and Skhirate. In the current work, we assess the involvement of *kdr* and *ace-1* mutations in the females adult *Cx. pipiens* mosquitoes from Mohammedia and Benslimane. The PCR diagnostic test was conducted by studying the sodium channel polymorphism in the two field populations and the AluI restriction site in the *ace-1* gene of the resistant individuals. After the double PCR, the three genotypes (Leu/Leu, Leu/Phe, Phe/Phe) were clearly identified in Benslimane region, but in Mohammedia, only two genotypes (Leu/Leu, Leu/Phe) were identified. The frequency of the resistant allele was strongly associated with the reduced mortality observed in bioassays with the WHO discriminating concentration of DDT in Benslimane region. This indicates that *kdr* is the major mechanism of organochlorines resistance in field populations in this area. The efficacy of organochlorines on resistant mosquitoes was shown to be dramatically reduced among mosquitoes from Benslimane region: 2% mortality after 24h exposure. It is not clear at this stage how the presence of *kdr* in the field will affect operational control. However, in view of the usual cross resistance conferred by this mechanism to other organochlorines and to pyrethroids, the efficacy of this groups of insecticides should be more detailed.

However, in Mohammedia region, the frequency of the resistant allele (F) wasn’t associated with the reduced mortality observed and 30% only of individuals carry the heterozygote genotype (Leu/Phe). Moreover, the resistant allele (Phe) in the heterozygote genotype could be the recessive allele and be not expressed in the phenotype nor influencing the resistance. But even *kdr* being recessive, it can also discriminate between heterozygotes and susceptible homozygotes (Martinez-Torres et al. 1998). Because the sodium channel region involved in *kdr* resistance is apparently highly conserved, the results obtained after the molecular diagnostic test could be used for monitoring pyrethroid *kdr* as part of the planning and implementation of vector control operations in these regions in Morocco.

In the second part, the PCR/RFLP diagnostic test that detects Two primers located in the third coding exon, on each side of the position 119 (Moustdir1 and Moustrev1) generated a 194 bp fragment by PCR on genomic DNA uncut by the AluI restriction enzyme either in resistant and susceptible mosquitoes from all regions. Therefore, the serine mutation conferring an AluI site was shown absent in all *Cx. pipiens* (N=20) resistant mosquitoes (homozygous and heterozygous) from Mohammedia and Benslimane and also in all the susceptible mosquitoes tested (homozygous susceptible, N=10).

Our results have demonstrated that all individuals either with only insensitive AChE (homozygous resistant), with only sensitive AChE (homozygous susceptible) or with both types of AChE (heterozygous) have none restriction site in all regions. The concordance between insensitive AChE and the presence of the G119S mutation, as diagnosed by the PCR test, was 0% (N=30) in all study sites. Therefore, the bendiocarb resistance observed in adults’ *Cx. pipiens* from Mohammedia and Benslimane certainly not due to AChE coded by *ace-1*. This situation suggests that are few other possible mechanisms of resistance that generate resistance towards OP or carbamate insecticides in *Cx. pipiens* in these regions.

Moreover, we already demonstrated that the pyrethroids resistance within mosquitoes from Mohammedia is completely metabolic and contributed to full involvement of P450 monooxygenases in the resistant populations (Aboulfadl et al. 2020). This finding aligns with our results that shown the absence of L1014F mutation within the same population from the same region. Otherwise, the resistance gained against DDT could be possibly due to DDTase. Contrary to pyrethroid-resistant *Cx. pipiens* from Benslimane that shown a limited involvement of CYP450 monooxygenases against pyrethroids (Aboulfadl et al. 2020). This observation confirms our result that demonstrated the presence of *kdr* gene in this population. The L1014F-*kdr* mutation confers the resistance towards both of pyrethroids and organochlorines. In the case of the cross-resistance to pyrethroids and organochlorines, the mutation L1014F affects the structure of the enzyme so that the catalytic site become less accessible to the insecticide. Consequently, the affinity between membrane proteins of neurons and insecticides decreases, which at the molecular level results in a mutation of the gene encoding the IIS4-IIS6 domain of the sodium channel. The *kdr* mutation is originated from a substitution of leucine by phenylalanine at position 1014 of the *kdr* gene. This mutation, common in West African *Anopheles gambiae* populations, confers a high level of resistance to permethrin and DDT, as well as cross-resistance to all pyrethroids (Martinez-Torres et al., 1998). For *Cx. quinquefasciatus*, the substitution of leucine – phenylalanine is commonly encountered in populations originating in North America and Africa. However, another mutation of the substitution leucine – serine (leu-ser), was observed equally and more frequently expressed in *Cx. quinquefasciatus* originated from Asia (Darriet, 2007).

The nature of the *kdr* gene in *Aedes aegypti* is both more complex and more diverse than for the genera *Anopheles* and *Culex* with the existence of five mutations, all different from the substitution’s leucine – phenylalanine and leucine – serine. The sequence of the gene encoding the sodium channel of this resistant population showed the existence of a Valine-Isoleucine mutation (Brengues et al., 2003). This resistance to pyrethroids remains a phenomenon whose extent is too often underestimated.

In Morocco, this mutation of the substitution leucine – phenylalanine at 1014 position was frequently observed in *kdr* gene in *Culex pipiens* populations in several regions: in Casablanca, Tanger and Marrakech (Bkhache et al. 2016), and now in Benslimane in the current study, but was proved absent in Mohammedia (Tmimi et al. 2018). The same mutation was observed absent in *Cx. pipiens* populations from Tunisia (Tabbabi et al. 2018).

Regarding *ace-1*^*R*^, the mutation results from a substitution of glycine by leucine at position 119 of the *ace1* gene. It confers cross-resistance to carbamates and organophosphates. On *Cx. pipiens*, it was observed that high concentrations of organophosphate or carbamate insecticides, concentrations approaching the solubility limit, had no longer any effect on mutated acetylcholinesterase of this mosquito (Raymond et al., 1985). In plus, biochemical tests on two populations of *An. gambiae* from Côte d’Ivoire found resistant to carbosulfan also showed the presence of an insensitive acethylcholinesterase (N’Guessan et al., 2003). In a case of resistance present but absence of target site modifications, insensitive acetylcholinesterase specifically, other resistance mechanisms could be in play. Our study shows the absence of G119S *ace-1* mutation in all the *Cx. pipiens* populations from both regions Mohammedia and Benslimane despite existing resistance. This means that the resistance happened in these regions is most likely due to an overexpression of a specific detoxification enzymes. Faced with the absence of recommended bioassays or validated synergists to make the point on the involvement of esterases and GST enzymes, the expression of enzymes by RNA extraction remains the only way to detect the enzymes involved.

Apart from resistance to insecticides that affect the structure of GABA receptors, other resistance mechanisms encountered in insects are essentially generated by three groups of enzymes: gluthation-S transferases, esterases and oxidases.

Mutated GABA receptors are characterized by lower affinity of their sites against dieldrin and lindane (halogenated organochlorines). This type of resistance is all the more important to consider as new molecules such as fipronil (phenylpyrazoles) act on these sites and already no longer show the slightest efficacy on resistant GABA mosquito populations (dieldrin / fipronil cross-resistance). Glutahion-S-transferase (GST) enzymes, in their turn, are synthesized by the insect and promote enzyme-insecticide conjugation that produces fewer toxic metabolites. Esterases or hydrolases degrade ester groups into alcohols and acids. Since pyrethroids, carbamates and organophosphates have esters, esterases play a major role in their degradation. Oxidases or monooxygenases induce oxidation reactions that lead to the detoxification of insecticides. Although they have a particular affinity for pyrethroids, these enzymes degrade virtually all chemical families.

Our result highlights the evidence of resistance mechanism linked to target site modification on *Culex pipiens* collected from two regions from central Morocco. This study is agreed with the previous work published by Aboulfadl et al. (2020) for the same populations. No G119 *ace-1*^*R*^ was proved present in all the populations and only L1014F *kdr* has been found in Benslimane populations but not in mosquitoes from Mohammedia. The distribution of the allelic frequencies of the resistant allele and the type of the resistance mechanism varies according to the treatment method and the insecticide used.

## Conclusion

In this study we investigated the resistance mechanisms related to target site modification in two populations of *Cx. pipiens* collected from two regions in central Morocco. None of the mutation linked to the acetylcholinesterase protein coded by *ace-1* gene has been found in all the populations. However, the mutation linked to the voltage gated sodium channel (vgsc) coded by *kdr* gene: L1014F has been found in only the resistant populations from Benslimane but not in the resistant mosquitoes from Mohammedia. Hence, the results obtained by the molecular diagnostic tests are in good agreement with the bioassay data published previously (Aboulfadl et al. 2020). This result gives important informations about the existing mutations on *Culex pipiens* from Benslimane and Mohammedia. This data could help authorities to better manage the resistance insecticide within mosquito populations.

